# Discovery of neuronal cell types by pairing whole cell reconstructions with RNA expression profiles

**DOI:** 10.1101/2024.12.30.630829

**Authors:** Tiago A. Ferreira, Mark Eddison, Monique Copeland, Mary L. Lay, Monet Weldon, Judith Baka, Emily Tenshaw, Alyssa K. Stark, David Schauder, Donald J. Olbris, Konrad Rokicki, Nelson Spruston, Paul W. Tillberg, Jayaram Chandrashekar, Wyatt Korff, Joshua T. Dudman, MouseLight Project Team

**Author notes:** Allen Institute for Neural Dynamics, Seattle, WA, USA.

## Abstract

Effective classification of neuronal cell types requires both molecular and morphological descriptors to be collected *in situ* at single-cell resolution. However, current spatial transcriptomics techniques are not compatible with imaging workflows that successfully reconstruct the morphology of complete axonal projections. Here, we introduce a new method, which we call morphoFISH, that combines tissue clearing, submicron whole-brain two photon imaging, and Expansion-Assisted Iterative Fluorescence *In Situ* Hybridization (EASI-FISH) to assign molecular identities to fully reconstructed neurons in the mouse brain. We used morphoFISH to molecularly identify a previously unknown population of cingulate neurons projecting ipsilaterally to the dorsal striatum and contralaterally to higher-order thalamus. By pairing whole-brain morphometry, improved techniques for nucleic acid preservation and spatial gene expression, morphoFISH offers a quantitative solution for discovery of multimodal cell types and complements existing techniques for characterization of increasingly fine-grained cellular heterogeneity in brain circuits.

## Introduction

There is broad agreement that a critical step to understanding the remarkable complexity of the mammalian brain is to develop an increasingly detailed and comprehensive understanding of its molecular constituent cell types and their complex projection patterns^1^. The ability to ultimately take such parts lists of cell types and assemble them into increasingly sophisticated models of brain function will depend both on understanding how molecular diversity contributes to the development and physiology of neurons as well as understanding the detailed connectivity of the diverse neuronal types. Connectivity is determined in substantial part by the detailed morphology of long-range axons that can span the entire mammalian brain ^2–4^. For example, the elaboration of cortex over mammalian development is substantially driven by the elaboration of subtle morphological and molecular variants of long-range projection neurons that are thought to be critical to empower new functions ^5^.

There has been remarkable progress over the last several years on complementary fronts: complete reconstructions of the extended morphology of neurons throughout the entire brain^3,4,6–8^ and increasingly deep analysis of the molecular constituents of individual cells *in situ*^6,9,10^. However, for a host of technical reasons, these advances have also made these two major classes of measurement incommensurate. Thus, to date, it has not been possible to obtain complete anatomical reconstructions and molecular cell-type characterization from individual cells in the same tissue. Rather, it has generally been necessary to compare across datasets to identify average projection patterns of molecularly defined cell types ^2^ or infer the likely molecular profile of individual reconstructed neurons ^11,12^. Pioneering approaches to spatial transcriptomics and single-cell barcoding have produced great insight into the intersection of whole brain projection targets and molecular profiling^4^. However, a critical unmet need is the development of methods that allow one to combine complete and detailed morphological reconstruction at submicron resolution with molecular characterization in individual neurons.

To accomplish this goal, we sought to develop a pipeline that combines molecular identification and complete morphological reconstruction of the same neurons. This required refining software to obtain complete and accurate single neuron morphological reconstruction, as well as developing a new method that preserves ribonucleic acids (RNA) and allows for multi-round fluorescent *in situ* hybridization (FISH) after weeks of processing time required for whole brain imaging. To obtain the highest topological fidelity, we quantified the most common pitfalls to neuronal tracing and used that data to refine workflows and software that obtain complete, single neuron reconstructions in a high-fidelity regime. To permit molecular profiling of these neuronal types *in situ*, we built upon the EASI-FISH approach to develop a protocol with vastly improved methods for preservation of RNA many weeks after fixation, tissue clearing, and imaging.

Here we describe a substantial addition of hundreds of highly accurate, complete neuronal reconstructions distributed across multiple cell types in the mouse brain, spanning a range of reconstruction problems from massive branching in midbrain dopamine neurons to long axons with many sparse branches of neurons in the in multiple areas along the neuraxis from neocortex to the pontine brainstem. Moreover, we describe and share HortaCloud, a software platform that greatly facilitates multi-user tracing in the cloud and validation of reconstruction quality. As a proof of principle, we describe the use of morphoFISH for the discovery of a rare and previously undescribed morpho-molecular subtype of long-range projection neuron specific to the cingulate cortex in mice.

## Results

### Release of new morphological reconstructions

The complete reconstruction of single axons in mammalian neurons is both essential and profoundly challenging^13^. A significant constraint on the function of a neuron are the other brain areas and cell types to which it connects via its axonal projections. In the previous data releases for the MouseLight Project Team ^3^, we have observed a broad distribution of complete axonal length and number of brain areas targeted from a sampling of ∼1200 reconstructed neurons across multiple, but necessarily limited cell type and brain area sampling. Here, we release an additional 536 fully and accurately reconstructed neurons. These include a subset of neurons from frontal cortical areas that are representative of the subset of neurons with the longest axonal arbors relative to other neurons (**Fig. 1a**). This allowed direct, quantitative comparison to recent large datasets that focused on the same populations of frontal cortical neurons^7^. In addition, we now include reconstructions from underrepresented subcortical areas including neurons from the midbrain and hindbrain and a sparse set of reconstructions from rare cell types that provide exemplars defining the bounds of the overall distribution observed across all reconstructions to date (**Fig. 1a** and **Data Availability**).

**Figure 1.**
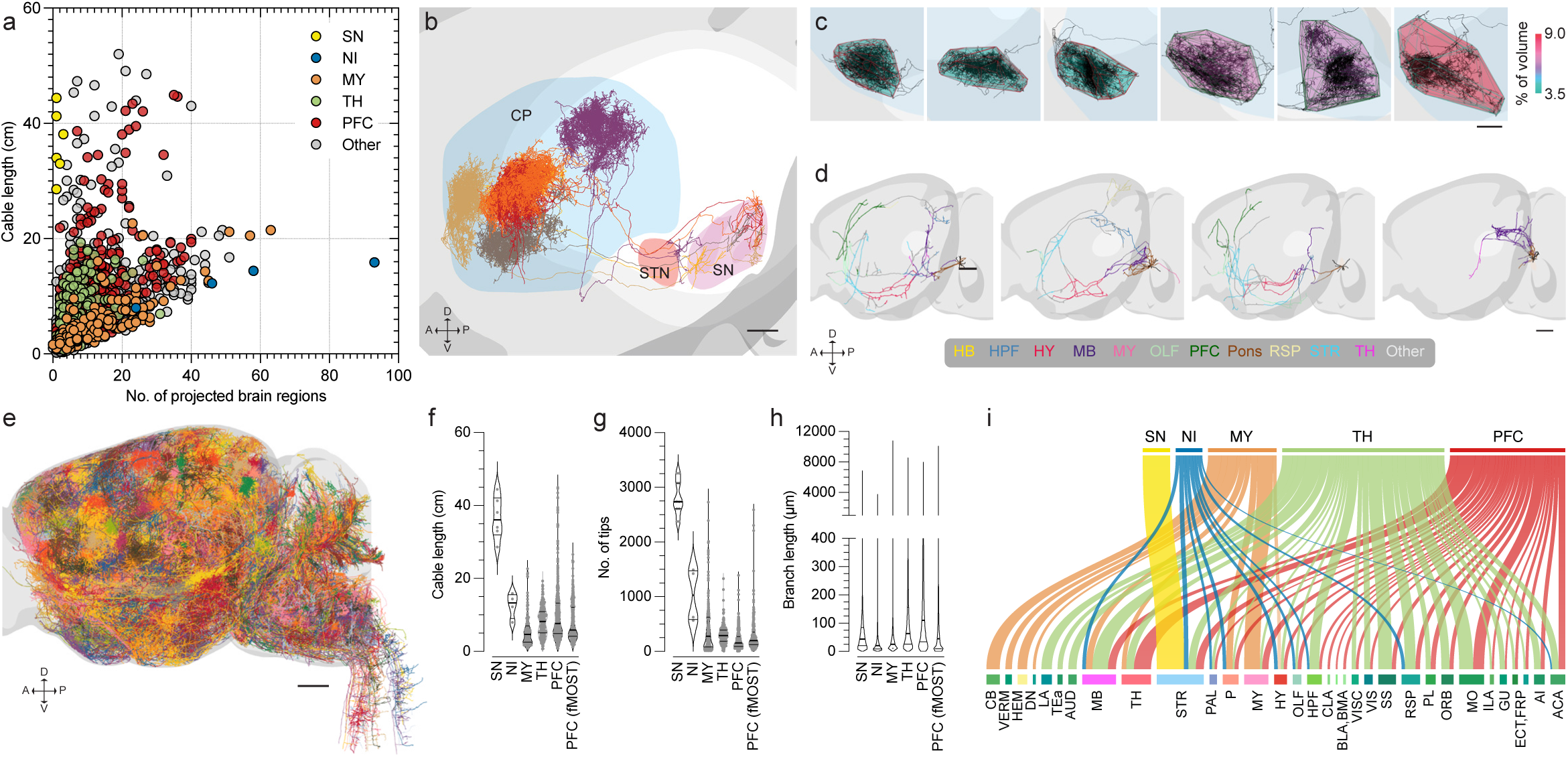
Whole-brain sampling reveals broader morphological diversity of neuronal projection types. **a.** The mouse brain (Allen Mouse Brain Common Coordinate Framework (CCF) was divided into 141 mid-ontology regions and cable length of individual axons plotted against the total number of unique regions innervated by their axonal endings (N=1638 cells). Cells from Substantia Nigra (SN, N=6), Nucleus Incertus (NI, N=4), Medulla (MY, N=142), Thalamus (TH, N=206), and prefrontal cortex (PFC, N=420) are indicated. PFC is defined by Anterior cingulate area (ACA), Infralimbic area (ILA), and Prelimbic area (PL). **b–d**. Detailed analysis of populations highlighted in **a**. dorsoventral (D-V) and anterior-posterior (A-P) axes are indicated. **b–c**. SN neurons projecting exclusively to the ipsilateral caudoputamen (CP). Single axonal tufts can cover as much as 9% of the ipsilateral CP lobe, as measured by the 3D convex hull formed by the axon end points. Subthalamic nucleus (STN) is also highlighted. Scale bars: 500µm. **d.** NI neurons in the pontine brainstem projecting to many areas in the forebrain (including hippocampal formation (HPF), hypothalamus (HY), olfactory areas (OLF), PFC, striatum (STR), and TH), retrosplenial area (RSP), midbrain (MB), and hindbrain (HB, including Pons (P) and Medulla (MY)). Scale bar: 1mm. **e.** Overview of new reconstructions included in the data release accompanying this manuscript (N=553). Scale bar: 1mm **f–h.** Morphometric properties of the 5 groups of cells described in **a**. PFC reconstructions (ACA, ILA, and PL) from Gao L. et al. 2022 (fMOST, N=426) are included for comparison. **i.** Sankey diagram summarizing axonal projection patterns. Flow weights reflect normalized innervation to target areas (see Methods). Leaf abbreviations reflect CCF nomenclature. Spinal cord projections were excluded from analysis. N=6 (SN); 3 (NI); 142 (MY); 206 (TH); 420 (PFC).

Cable lengths of individual neuron axonal arbors in the mouse brain can range from a few millimeters to total up to ∼50 cm long^3^. These longest cable lengths can be observed in neurons that connect very densely to just a single brain region by elaborating complex and plentiful short terminal axonal branches. This edge of the distribution is typified by midbrain dopamine neurons of the substantia nigra (SN; **Fig. 1a–c**). At the other extreme, a neuron may elaborate a similar cable length of axon and yet innervate several tens of different brain regions. This is typified by projection neurons of the nucleus incertus (NI; **Fig. 1a,d**). Cell types from the forebrain (e.g. cortex), midbrain (e.g. thalamus), and hindbrain (e.g. medulla, NI) fill in an approximately continuous distribution between these extremes and are included in the new data release (**Fig. 1e**).

Despite very large differences in total cable length and number of terminal points (‘tips’) across populations, the branch length distributions are notably similar (**Fig. 1f-h**). This broad range of axonal complexity presents a number of critical challenges for reconstruction. In particular, it is often the case that a neuron may have a single or few axonal branch points located somewhere along centimeters of axon that determines its connectivity to one or several brain areas via characteristic variation in the number of terminal branches (**Fig. 1g**) and shared, but very broad distribution of axonal branch lengths characteristic of all neuronal types examined to date (**Fig. 1h**). Accurate identification of all brain areas targeted by single neurons is a critical functional feature and essential for identifying morphotypes of projection neurons and distinguishing amongst related projection neurons from a given area revealed by observing the distribution of target innervation areas across the neuraxis from the forebrain to hindbrain **(Fig. 1i)**. Neurons that do not innervate a subset of targets known from a given area is an important quantitative difference for the evaluation of function and critically depends upon the reconstruction method used and the relative certainty about completeness and accuracy.

### Development of morphoFISH

Complete axonal tracing in mouse brain tissue requires extensive chemical modification of the brain tissue to accommodate whole brain imaging (**Fig. 2a**). This has been previously achieved at the expense of RNA integrity. Indeed, RNA is barely detected in MouseLight imaged tissue^3^ (**Fig. 2b**). Taking inspiration from EASI-FISH^14^—a highly sensitive technique that stabilizes RNA by capturing it in an expandable hydrogel—we developed a protocol that prevents RNA degradation during histological handling and imaging of the brain by embedding in a non-swelling polyacrylamide hydrogel. After whole brain imaging, regions containing cell somata are later targeted for expansion and EASI-FISH processing. The method both increases the initial RNA retention and slows down its degradation. We took a number of steps to preserve RNA in whole-brain preparations. First, we implemented a 3-step transcardiac perfusion in which the animal is first perfused with paraformaldehyde (PFA) alone; then with methacrylic acid-NHS ester, which reacts with proteins and will later anchor them to the gel; then with a solution including acrylamide monomer, bis(acryloyl)cystamine crosslinker, and the APS/TEMED initiator system, which will later polymerize to form the gel. A polymerization inhibitor (4-hydroxy-TEMPO) is added to this mixture to ensure gelation solution has time to diffuse throughout the brain before the polymerization reaction takes place. Once the brain is dissected and gel allowed to polymerize, it becomes embedded in an acrylamide gel where proteins are covalently anchored to the gel polymer (**Fig. 2a**). PFA post-fixation is complemented with glutaraldehyde (GA) and melphalan, an alkylating agent that reacts with nucleic acids and bears a primary amine that is fixable with aldehydes^14,15^, creating extensive crosslinking amongst proteins, RNA, and the acrylamide gel. Indeed, with this protocol, we effectively doubled the initial capture of total RNA in the brain when compared to prior perfusion approaches (**Fig. 2b**).

**Figure 2.**
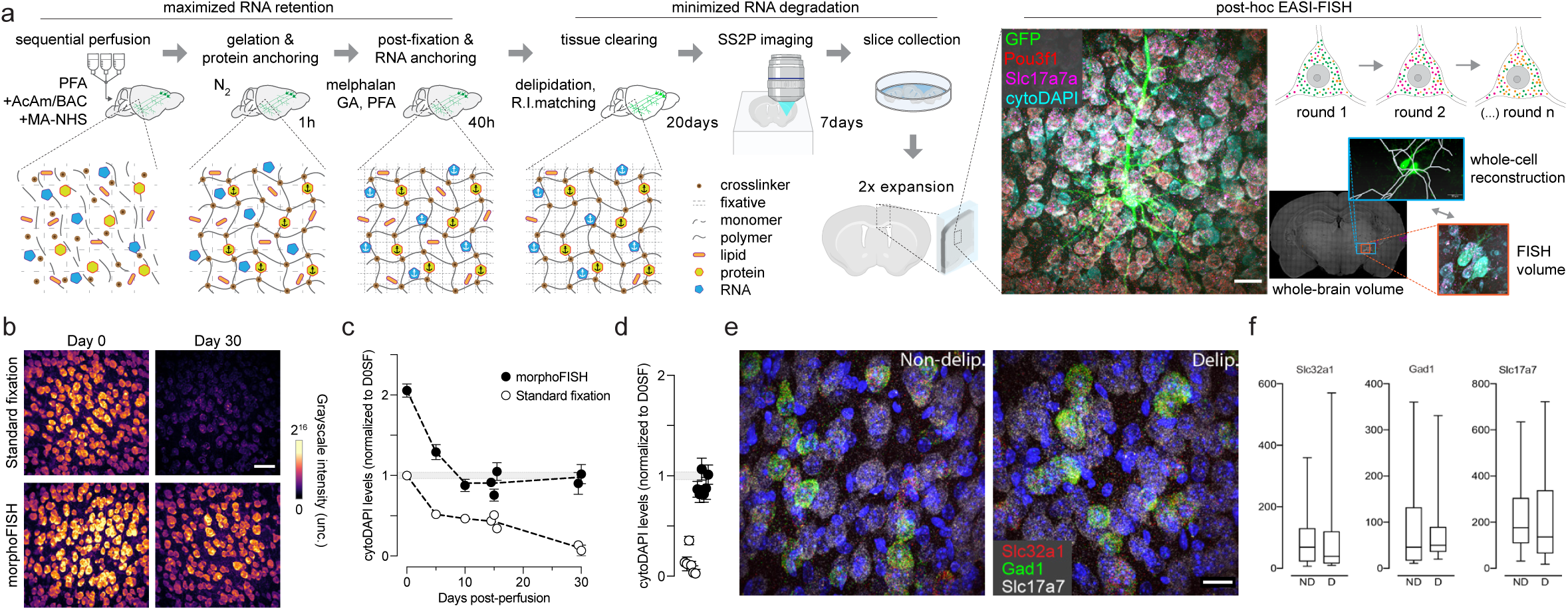
morphoFISH allows for sensitive detection of transcripts in cleared, imaged tissue. **a**. Diagram of the morphoFISH protocol. Paraformaldehyde (PFA), acrylamide (AcAm) monomers, AcAm cross linker (BAC), and amine-reactive MA-NHS are sequentially perfused into a brain with sparsely labeled neurons expressing a fluorescent protein (FP). After brain dissection gel is polymerized and proteins are covalently anchored to the gel. During the post-fixation step, PFA fixation is complemented with glutaraldehyde (GA), and RNA is cross-linked to the scaffold of gel/proteins by the alkylating agent melphalan. Inserts depict a conceptual diagram of RNA anchoring to the hydrogel scaffold within the fixed tissue. Next, the whole brain is cleared by removal of lipids and index-matched to a high refractive index solution, and imaged on a serial-sectioning two-photon (SS2P) microscope. Duration of individual steps is indicated. After imaging, sections are harvested and multiple rounds of EASI-FISH performed on expanded sections with FP positive somata and imaged by light-sheet microscopy. Preservation of FP signal during EASI-FISH enables labeled neurons to be mapped back to their original location in the SS2P whole brain volume, and assigned to its morphological reconstruction. **b,c**. Cytosolic DAPI-stained RNA (cytoDAPI) monitored in 200µm-thick brain slices left at room temperature for 30 days. **b.** Representative images **c.** Time-course quantification of cytoDAPI signal. morphoFISH allows for a two-fold detection of RNA relatively to Day 0 standard fixation protocols (D0SF). While loss of RNA occurs in both conditions, the high-retention provided by morphoFISH yields comparable D0SF levels at day 30, while RNA is virtually undetected under standard fixation. Individual data points reflect measurements from 300-500 cells per slice (mean±SD). Dashed line connects means between sampled days. **d**. Quantification of cytoDAPI levels after a mock imaging experiment in which cleared tissue (delipidated and R.I matched) was left on the imaging tray of the SS2P imaging rig for 5 days. **e,f**. Impact of delipidation on morphoFISH transcript detection for Slc32a1, Gad1, and Slc17a7 (counts/cell). **e**. Representative images (nuclei labeled in blue) **f**. Spot quantification for segmented cells in **e**. Scale bars: 50µm.

While increasing initial capture was a critical first step, but subsequent degradation was an even more critical issue that led to almost no RNA preservation in standard approaches (**Fig. 2b**). We next focused on mitigating RNA loss during the weeks-long histological processing of the tissue, which includes delipidation and refractive index (RI) matching. While the former is aimed at removal of light-scattering lipids from the tissue, the latter is aimed at minimizing the spatial variation of refractive index due to the heterogeneity of tissue components. Combined, these are the longest processing steps and proved to be specifically problematic for RNA preservation in tissue. To monitor detection sensitivity after removal of lipids, we compared tissue with and without delipidation for abundant RNA transcripts. We found similar expression levels across individual cells regardless of whether delipidation had been performed (**Fig. 2e,f**). We then tested refractive index matching strategies ^16–19^ for RNA retention and settled on the DMSO/iodixanol cocktail already adopted by our platform ^3,20^. To counteract RNA degradation during refractive index matching and imaging processes we supplemented the imaging bath with polyvinylsulfonic acid (PVSA), a negatively charged polymer with side structures mimicking the conformation of mRNA molecules, that has been shown to be an effective competitive inhibitor for ribonucleases^21^.

In total, the complete morphoFISH protocol was successful at yielding approximately the same preservation of RNA after 30 days post-perfusion as achieved with standard fixation procedures in prior EASI-FISH protocols (day 0 standard fixation, D0SF) (**Fig. 2b,c**). Consistent with improved RNA retention, cleared tissue left unattended for 5 days in the serial-sectioning two-photon microscope used by MouseLight retained comparable D0SF levels (**Fig. 2d**). Typically, reconstruction of complete axonal arbors requires signal amplification of fluorophores along dim axon collaterals using whole brain immunohistochemistry ^3^. This procedure would take several weeks on top of the 30 days to process the brain and is therefore prone to compounded mRNA loss. Rather than incorporating this extra step into morphoFISH, we focused on optimizing a pipeline for reconstructing single neurons labeled with endogenous, unamplified fluorescent reporters, which enables effective, albeit somewhat lower yield labeling^22^.

### High-fidelity neuronal reconstructions

We obtained sparsely labeled neuronal populations by injecting a mixture of low-titer AAV Cre and a high-titer Cre-dependent reporter as previously described^3^. In order to further improve reconstruction accuracy, we first quantified the most common errors afflicting reconstructions of long-range axonal projections. Second, we used insights from the most problematic errors to optimize the development of our custom reconstruction software, HortaCloud, in preparation for making it available to the broader scientific community. Third, we released a standardized protocol and guidelines ^23^ for tracing long-range projections using the open HortaCloud ^24^ tool (**Fig. 3**).

**Figure 3.**
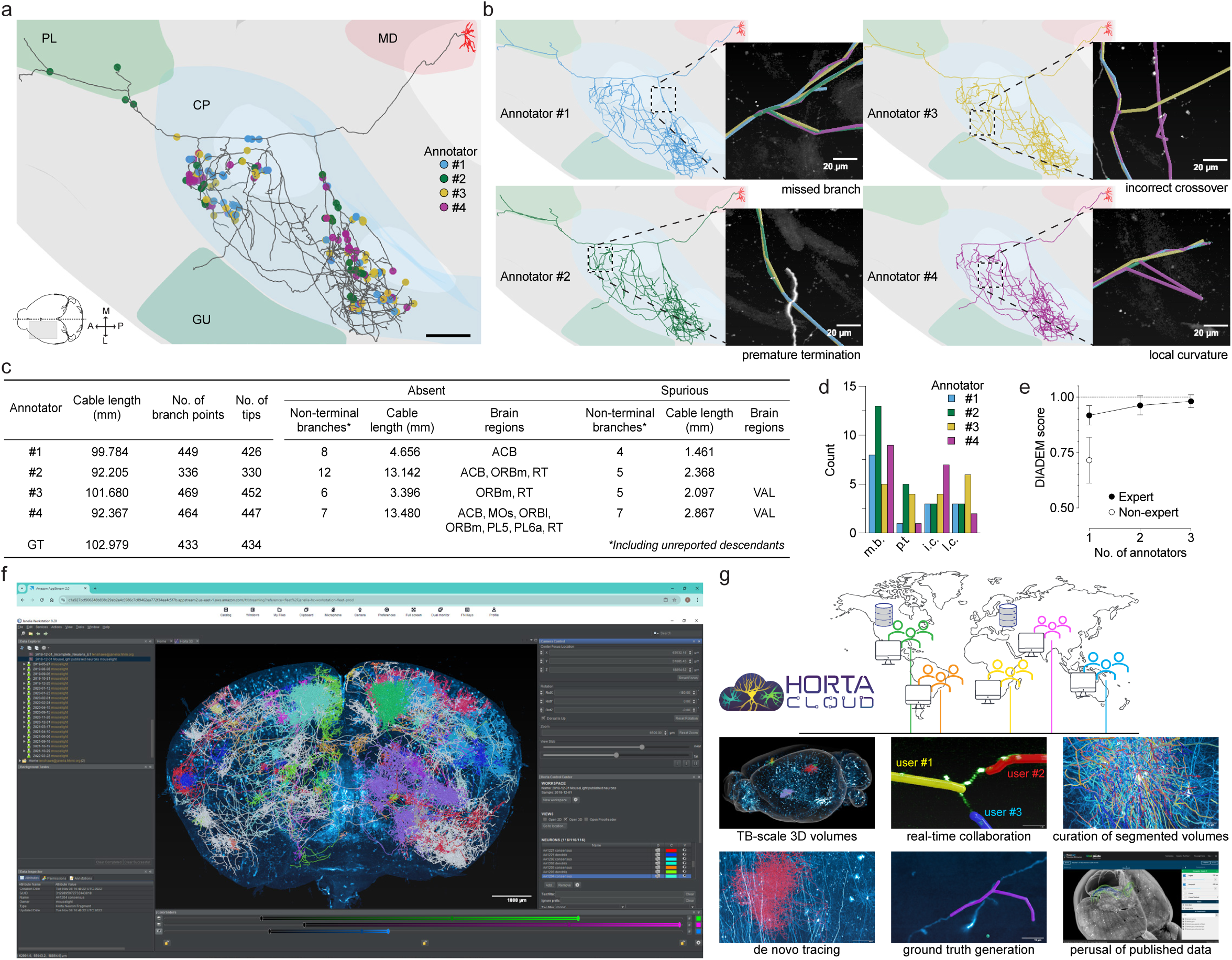
Axonal reconstructions guided by quantitative indicators of rigor. **a–d**. The axonal arbor of an ipsilateral mediodorsal thalamic neuron (MouseLight ID AA1624) was independently reconstructed by multiple expert annotators and tracing errors quantified. **a.** Ground-truth (GT) obtained from the four individual reconstructions. Markers highlight the locations at which major errors occurred. Note that when subsidiary branches are involved, only the first error in the parental branch is marked. Prelimbic area (PL), Caudoputamen (CP), Mediodorsal nucleus of thalamus (MD), and gustatory areas (GU) are highlighted. Dendrites are displayed in red and were excluded from analysis. Bottom-left diagram highlights the location of the neuron in the mouse brain. Midline (dashed line) and medio-lateral/anterior-posterior axes (M-L/A-P) are indicated. Scale bar: 500µm. **b.** Result of individual annotators. Inserts highlight the four most common types of detected errors (see Methods). **c.** Morphometric properties of axonal reconstructions in **a,b**. Table includes sections that were not traced despite being present in GT (“Absent”), and sections from neighboring cells that were traced but do not exist in GT (“Spurious”). Brain regions (as defined in Fig. 1a) associated with at least one axonal termination are listed. Note that non-terminal branches typically ramify into several subsidiaries containing downstream errors not included in this analysis. **d.** Absolute frequencies of categorized errors grouped by annotator: missed branch (m.b.), premature termination (p.t.), incorrect crossover (i.c.), and local curvature (l.c.). **e.** Increase in accuracy as a function of the number of expert annotators converging into GT, as assessed by pairwise DIADEM scores (mean±SD) against the GT consensus. For context, scores from 12 non-expert annotators are also included. Scores were computed using default settings without adjustment of parameters. **f.** Screenshot of the HortaCloud user interface running in a web browser. Quantitative error analysis guided its development to favor reconstruction accuracy. **g.** Conceptual illustration of HortaCloud’s architecture. Centralized servers (“instances”) host shared workspaces (directories that aggregate 3D volumes, metadata, and annotations) that can be accessed concurrently by multiple users (“clients”) from internet connected locations. The ability of users to access and modify workspaces is managed by a permissions’ framework. Vignettes highlight key system capabilities.

For error quantification, expert annotators independently reconstructed the same cell, and a ground-truth reconstruction was generated by comparing their work (**Fig. 3, and Methods**). After repeated tracing of a mediodorsal (MD) thalamic neuron with more than 100 mm of cable length, we found that: 1) tracing errors could be grouped into four categories; 2) errors were spread across the ramified sections of the axonal arbor; 3) errors were common even amongst expert tracers; and 4) errors occurred at frequencies that were low but nonetheless sufficient to distort the topology of the cell and complete annotation of its target areas (**Fig. 3a–d**). We found that ‘crossovers’ between neurites (either from the same cell or from neighboring cells) have the highest potential to alter projection patterns while being the most time-consuming to resolve. For many neuronal morphotypes, this can often require expert perusal of as much as extra 12cm of cable length per cell, because it may involve tracing neighboring neurites from multiple cells (**Fig. S1**). Building on previous work^3^, we found that assembling consensus reconstructions from two independent annotators provides an acceptable balance between accuracy and throughput. However, tracings obtained from a large group of annotators that did not follow expert guidelines were found to be of low quality, as assessed by DIADEM score comparisons^25^ (**Fig. 3e**). Collectively, these results are at odds with previous studies proposing that spot-checks after automated segmentation are sufficient to obtain acceptable accuracy^7,26^. We found that spot-check approaches tend to be biased towards bright neurites, however, fluorophore availability (brightness) is not strongly correlated with the ability to fully reconstruct long-range projections (**Fig. S2**). This may contribute to the observed quantitative differences in the distribution of cable lengths between MouseLight datasets and reconstructions from frontal cortical neurons by other groups^7^ (**Fig. 1f-h**).

### Identification of a novel morpho-molecular neuron subtype

The successful preservation of RNA in tissue allows multi-round *in situ* hybridization-based molecular profiling of individual neurons within tissue suitable for complete tracing of sparsely labeled neurons. While this approach is necessarily limited in throughput, it can nonetheless, in principle, empower the molecular profiling of neurons with unique axonal morphologies.

Areas of the frontal cortex, in particular cingulate areas, harbor molecular subtypes of deep layer, putative long-range projection neurons that are unique and not found in other cortical areas^12^. Despite a detailed molecular characterization of the populations of neurons in frontal areas, it is currently not possible to map these rare or unique molecular subtypes onto unique morphological projection patterns, which will likely be critical for complete cell typing ^1^. Building upon our extensive tracing of individual neurons in frontal cortical areas we looked for individual neurons with rare, previously underappreciated axonal projection patterns. We identified one particular neuron with a highly unique projection pattern based upon previous examinations of bulk and large scale single neuron reconstructions. Specifically, we identified a neuron in the cingulate cortex that projects ipsilaterally to the dorsal striatum (STR or the caudoputamen, CP) and contralaterally to the higher-order parafascicular thalamic nucleus (PF) (**Fig. 4a–c**). This projection pattern, for example, was never observed in extensive reconstructions previously published in anterior lateral and primary motor areas ^3^. Given the absence of any such combination of projections in a single neuron in other cortical areas, this suggested that this neuron could represent a unique morphomolecular subtype.

**Figure 4.**
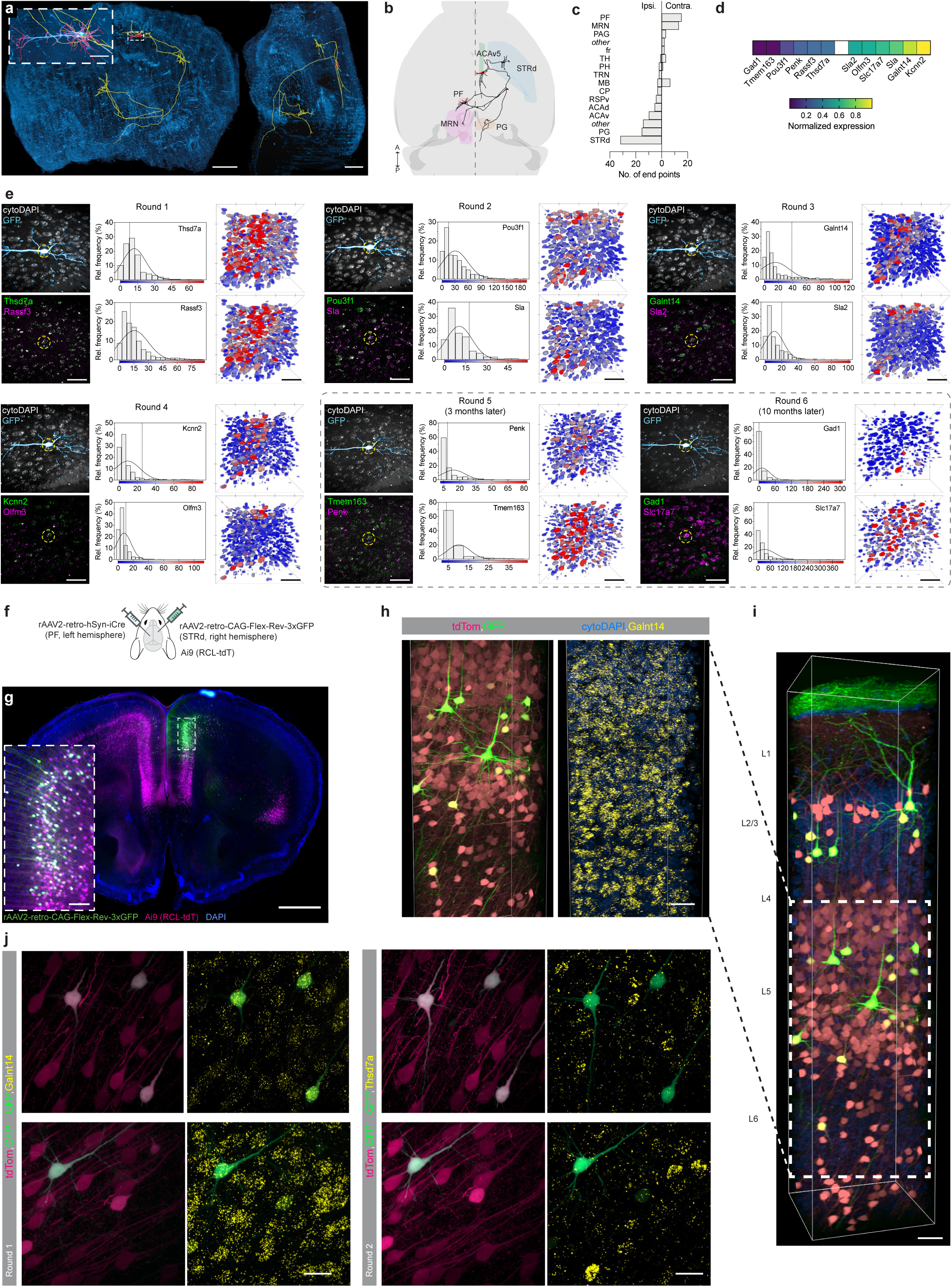
Molecular characterization of a novel class of cingulate neurons. **a.** Coronal (left) and sagittal (right) views of a ventral anterior cingulate area layer 5 (ACAv5) neuron (AA1624) traced in HortaCloud. Dendrites are magnified in insert and highlighted in red, with axon in yellow. Scale bars: 1mm (whole brain), 100µm (insert). **b.** Dorsal view of the same cell in CCF coordinates, depicting ipsilateral projections to the dorsal striatum (STRd) and pontine gray (PG), and contralateral projections to the parafascicular nucleus (PF) and the midbrain reticular nucleus (MRN). Dashed line highlights midline. Anterior-posterior (A–P) axis is indicated. **c.** Frequency histogram of end points of the same neuron grouped by brain areas and split by ipsilateral/contralateral hemisphere. “Other” aggregates brain areas associated with a single end point or non-cellular regions. Abbreviations according to CCF nomenclature: ACAd: Anterior cingulate area dorsal part; fr: fasciculus retroflexus; MB: Midbrain; MRN: Midbrain reticular nucleus; PAG: Periaqueductal gray; PH: Posterior hypothalamic nucleus; TH: Thalamus; TRN: Tegmental reticular nucleus. **d.** Summary of the gene-expression profile of AA1624 (detailed in **e**). Reported values are normalized to the 10^th^ and 90^th^ percentile of spot counts for all cells probed. **e.** morphoFISH characterization of AA1624 (12 transcripts probed across 6 rounds). For each round, a field of view centered on the cell’s soma was imaged for cytoDAPI, GFP, and two HCR-probed transcripts. A maximum intensity Z-projection (depth: 100µm) is shown. Histograms depict counts of HCR spots in cytoDAPI-segmented somata. Vertical line highlights the position of the cell in the distribution. Overlaid gaussian fit reports mean and standard deviation. Heatmaps depict cytoDAPI-segmented somata color-coded by the number of HCR spot counts scaled to the lowest and highest count detected in the volume. To demonstrate robustness of detection, the last two rounds were performed several months after round 1, as indicated. **f.** Diagram of dual-stereotaxic injection, in which an Ai9 (RCL-tdT) reporter mouse was injected with rAAv2-retro-CAG-Flex-Rev-3xGFP in STRd (right hemisphere), and AAv2-retro-hSyn-iCre in PF (left hemisphere). **g.** Expression pattern of dual-stereotaxic injection in the PFC (see **Fig. S3**, **S4** for complete expression pattern). Magnified insets depict somata of STRd^IPSI^/PF^CONTRA^ GFP^+^/tdTom^+^ cells in the ACA. Scale bars: 1mm and 100µm (inset). **h–j.** Molecular characterization of GFP^+^/tdTom^+^ cells in ACAv5 described in **f**,**g**. **h**. Expression of Galnt14 in Layer (L) 5. **i**. Overview of cortical column. **j**. Magnified inserts depicting GFP^+^/tdTom^+^ cells expressing high levels of Galnt14 relative to Thsd7a. Scale bars: 100µm (i,j), 50µm (j).

Following complete tracing of this neuron, we were able to profile the molecular expression pattern through several rounds of non-destructive *in situ* hybridization using methods modified from the EASI-FISH protocol previously described^14,27^ (**Fig. 4d**). Taking advantage of critical resources of gene expression profiling using deep sequencing of cells in the cingulate cortex, we identified and validated a family of marker genes sufficient to identify unique molecular types of deep layer projection neurons (**Sup. Information**). Using multiple rounds, we probed the target neuron with 6 rounds (2 probes each) of *in situ* hybridization and imaging. Quantification of expression level of each gene across the population of neurons within the imaged tissue section identified the target neuron as positive for Galnt14 and Kcnn2 and negative for a host of other genes—consistent with a unique subclass of deep layer projection neuron found in cingulate cortex. This connects a unique morphotype of a projection neuron to relatively rare molecular subtypes of projection neurons in the cingulate/retrosplenial cortices identified from extensive sequencing of cortical neurons in the mouse brain^28^.

We next sought to confirm the molecular identity of projection neurons with this unique innervation pattern and discover the relative prevalence of this morphotype in a higher throughput manner. Identification of the unique projection properties allowed us to design an intersectional viral labeling scheme. We injected retrograde viruses into the contralateral PF (expressing Cre-recombinase) and ipsilateral dorsal striatum (expressing GFP) in 3 tdTomato cre-dependent reporter mice (**Fig. 4e**). To highlight the remarkable RNA retention achieved by this protocol we performed additional rounds of profiling at 3 months and 10 months after perfusion (**Fig. 4e**). While some loss of total RNA was apparent by 10 months there was still sufficient retention for the detection of abundant transcripts that can be critical for broad molecular classification.

The unique morphology of the target neuron should be reflected in a population of dual-labeled (tdTomato+ and GFP+) ipsilateral to the dorsal striatal injection site and located in the cingulate cortex. This is expected to be contrasted with a much larger population of tdTomato+ deep layer projection neurons ipsilateral to the Pf injection site, as thalamocortical projections are primarily ipsilateral and not derived from the bilateral intratelencephalic projection subtypes that innervate striatum^3^. To provide a comprehensive estimate of the relative prevalence of the STR^IPSI^/PF^CONTRA^ morphotypes, we performed mesoscale imaging of the majority of the rostral brain and confirmed that the intersectional labeling strategy did identify a small but clear population of STR^IPSI^/PF^CONTRA^ projection neurons localized to the rostral cingulate cortex (**Fig. 4f**). Finally, to confirm that this population was consistent with the morphomolecular type identified from morphoFISH, we performed FISH on the labeled tissue (**Fig. 4g**). We found that the dual-labeled subset of STR^IPSI^/PF^CONTRA^ projection neurons were indeed excitatory neurons (Slc17a7^+^) from the subset of Galnt14^+^ deep layer projection neurons in the cingulate.

## Discussion

Here we present several innovations, including refinements to techniques for tissue preparation and software, for efficient, complete, and accurate reconstructions that combine to empower simultaneous multi-round *in situ* hybridization-based molecular profiling of individual neurons with complete axonal reconstructions in the mouse brain—a resource we refer to as morphoFISH. We provide one particular example of a unique morphomolecular cell type identified in the cingulate cortex. Discovery of this unique cell type could be rapidly translated into customized, higher-throughput techniques suitable for future functional experiments. The cingulate cortex is thought to contain a number of unique molecular and presumed anatomical types of projection neurons that may contribute to the unique and critical functions of the cingulate cortex, two areas implicated in motor skill learning in rodents ^29,30^. The sharing of output mediated by these unique projection neurons to ipsilateral high-order thalamic areas and contralateral dorsal striatum may play critical roles in interhemispheric and interregional coordination critical for motor skill learning.

While morphoFISH presents a unique resource that can be a critical component for a broad effort to identify morphomolecular cell types in the mouse brain, it also presents some fundamental challenges to throughput. Molecular profiling via repeated rounds of *in situ* hybridization is powerful and has been used to discover novel functional correlates of molecularly defined cell types ^31^. The approaches for dramatically improved RNA retention described here are general and offer the promise of enhancing many approaches to molecular profiling, including the improved characterization of low-expressing genes that may be critical for some questions. However, while this approach offers potentially very high sensitivity for individual transcripts, it also requires substantial time due to modest (≤4) multiplexing as compared to other, higher throughput approaches for spatial transcriptomics. Autofluorescent lipofuscin granules start to accumulate in the mouse brain after 8 weeks ^32^, and so we set our quantification routines to ignore lipofuscin foci. As a consequence, we are likely undercounting HCR spots across probing rounds. We predict that the same type of experiments performed using younger animals would allow for more straightforward quantification.

The recent advent of approaches allowing the full reconstruction of sparsely labeled neurons in the mouse brain has already yielded large databases that include thousands of reconstructed neurons. At the same time, this is a small sampling relative to the entire mouse brain and is heavily biased towards select neuron types in specific brain areas. Automatic processing via modern methods and software continues to rapidly develop and has dramatically improved the throughput of reconstruction. However, there remains a critical role of human intervention, as even exceptionally rare errors—e.g., a missing branch—can produce large differences in reconstruction accuracy. There is no ground truth to date, and thus differences in average properties across different reconstruction methods, a small subset of which we describe here, will be critical to evaluate carefully as reconstructed neurons proliferate. In *Drosophila*, light-level reconstructions have been indispensable for interpreting EM connectomes and have guided retrieval of molecular, functional, and mesoscale data used to catalog neuron types in detail ^33^. Given the expert curation of our morphologies, and the multi-modal characterization enabled by morphoFISH, we predict such data will be critical to facilitate similar advances in the mouse. Moreover, the MouseLight database provides reconstructions of subcortical populations distributed throughout the neuraxis from forebrain to hindbrain that have not been studied in as much detail to date.

The detailed reconstructions and multi-user annotations that constitute the database of reconstructed neurons described here were all performed in HortaCloud – a new cloud-based tool for collaborative and accurate neuron reconstruction. Unlike traditional desktop applications, HortaCloud enables collaborative tracing across remote locations using a web browser. HortaCloud builds on the proven track record of MouseLight’s Horta desktop application^3^ with a cloud system that any user or institution can easily deploy to avoid the complex and costly set up previously required for handling terabyte-sized anatomical datasets with on-premise infrastructure. HortaCloud presents a freely available and open source application that is already ‘future proofed’ for new imaging modalities and file types as they become available and includes a number of critical analysis tools to lower the burden for early adopters (e.g., EASI-Tower, SNT, Python notebooks) ^24^. Finally, to facilitate adoption of the methods described here, we have shared detailed, step-by-step protocols^20^ that democratize access to this technology.

## Methods

### Animals

Wild-type C57BL/6 animals were obtained from Charles River Laboratories and cre lines from the Jackson Laboratory. Genotype details are listed in the metadata of reconstructed cells downloadable from ml-neuronbrowser.janelia.org (see Data Availability). Adult females (12-20 weeks) were used for all experiments and were group housed with sex-matched littermates. Mice had access to ad libitum food and water and were housed in an enriched environment. All experimental protocols were conducted according to the National Institutes of Health guidelines for animal research and were approved by the Institutional Animal Care and Use Committee at Howard Hughes Medical Institute, Janelia Research Campus (Protocol #14–115).)

### Viral Labeling

Stereotaxic dual injections of rAAV2-retro helpers^34^ (Addgene plasmid #81070; RRID:Addgene_81070) were performed as previously described for single axon tracing anatomy^3^. For labeling of the unique ACA projection type injection coordinates were as follows: AP: -2.2, ML: 0.7, DV: -3.6 (PF); AP: 0.9, ML: 1.5, DV: -2.5 (STR). High-titer (∼10^12^ GC/ml) viruses: CAG-Flex-REV-3xGFP (STR, 400nL), hSyn-iCre (PF, 200 nL), were obtained from the Janelia Research Campus Viral Services and diluted as necessary. Injections were made into ‘Ai9’ reporter mice (JAX: 007909) which express tdTomato upon exposure to cre recombinase.

### Imaging

Dual Whole brain: Whole brain imaging was performed as previously described^3^. To ensure RNAse activity would not accumulate in the imaging bath over time, a peristaltic pump was used to replenish imaging solution at a constant flow of 1ml/min. An Ambion™ RNaseAlert™ lab test kit (ThermoFisher) was used to monitor RNAse activity and solution replaced in case of detection. Imaging progressed as described previously^3,22^ but brain slices (180–200µm) were collected every 2 cuts using a sterile brush and placed in 24-well plates. Each slice was washed (once in RNAse-free water and twice with 70% ethanol) and placed in 70% ethanol at 4°C until probing^27^. Serial-sectioning: Following standard perfusion and protocols, dissected brains were embedded in 5% agarose, and coronal sections (100µm thick) were cut on a vibratome (Leica). Sections were mounted in Vectashield® hardset media with DAPI (Vector Laboratories) and imaged on a confocal slide scanner using the TissueFAXS platform (TissueGnostics).

### Image Processing

Stitching, segmentation of neurites, and brain registration against the Allen CCF were performed as previously described^3^. Soma intensities were manually retrieved in HortaCloud and normalized to background fluorescence for a surrounding area without neurites, somas, or other noticeable features. Slide scanner images were processed in Fiji^35^, subscribed to the following update sites: 3D ImageJ Suite^36^, IJPB (MorphoLibJ)^37^, Neuroanatomy^38^, and sciview^39^. Volumes were rendered using 3D Viewer^40^ and sciview.

### Morphometry

Morphometry comparisons, volume analysis, connectivity analyses, and reconstruction renderings were performed in SNTv4.9.9^38^ using the provided scripts (see Data Availability). Dendritic reconstructions were excluded from all quantifications. Anatomical groups were defined by soma location.

*Cable length vs projected brain regions analysis*: Only CCF compartments of ontology depth 7 with publicly available meshes were considered (141 compartments). For each axon, brain regions associated with at least two endpoints were tallied.

*Convex hull analysis*: were computed as the smallest three-dimensional convex closure containing the end-points of an axon. Volume ratios were obtained by normalizing the volume of the convex hull to the halved volume of bilateral CP.

*Summaries of axonal projection patterns*: Sankey diagrams were obtained using SNT built-in functions: Target brain regions (diagram leaves) were empirically chosen. For each cell, cable length within each target region was computed and divided by the axon’s total cable length (TCL). To improve diagram readability, regions associated with less than 5% of TCL were not tallied. Averages of these percentages were then obtained from all the axons in each group and used as flow weights. Width of cell groups (diagram sources) is affected by the number of flows in the group and is not strictly proportional to the no. of cells in the group.

*Quantification of errors*: Differences between reconstructions passes were collected as “diff SWCs” as detailed in the companion protocol^23^. Errors were manually sorted into categories and mapped into the consensus reconstruction using nearest-match coordinates. DIADEM metrics^25^ were computed using default parameters.

*Quantification of effort required for crossover resolution*: Amount of cable length required to solve ambiguous crossovers was retrieved using HortaCloud built-in functions by collecting all traced fragments used to solve topological ambiguities at selected locations (Fig. S1) and summing their length. Locations were selected prior to reconstruction.

### Tissue preparation

Detailed morphoFISH protocols are published at protocols.io^20^. In short: Transfected mice were anesthetized with an overdose of isoflurane and then transcardially perfused with the following solutions sequentially: 1) perfusion fixative (4% PFA); 2) protein anchoring solution (4% PFA, 20mM MA-NHS); 3) gel solution (4% PFA, 20% AcAm, 0.1% BAC, 0.2% TEMED, 0.01% 4-Hydroxy-TEMPO, 0.2% APS). Brains were extracted and gel allowed to polymerize for 1 hour in a nitrogen-saturated enclosure. Brains were then post-fixed (4% PFA, 0.05% glutaraldehyde, 10mM MOPs (pH 7.0), 1mg/ml melphalan) at 4°C 40 hour) and washed in PBS to remove all traces of fixative. Delipidation and index matching proceeded as detailed in the companion protocols^20,41^, the brain was embedded in 4% low melting point agarose, and imaging solution supplemented with 10mg/ml PVSA.

### Quantification of total RNA

200 µm-thick brain sections were sliced on a vibratome and collected into individual wells of a 24-well plate in RNAse-free PBS (Invitrogen). Slices were collected at the reported time points (Fig. 2) and processed for cytoDAPI detection as previously described^27^. Tissue was imaged on a light-sheet microscope (Zeiss), cells segmented with Cellpose3^42^ (“cyto3” model) and intensities of segmented ROIs measured in Fiji^35^. Intensities were normalized to the overall mean obtained from D0SF slices.

### EASI-FISH

In-situ hybridization was performed as previously described^14^ but with an improved gel recipe for gel robustness, detailed in the companion protocol^27^ with two genes probed per round. A probe library of 31 genes was defined using data from previous RNAseq experiments^12,43^ (Table S1). Quantification of HCR spots was performed as follows: Cellpose3^42^ was used for 3D segmentation of cytoplasmic DAPI signal. The default “cyto3” model was used. Rounds 6 and 7 performed after a delay of several months featured low cytoDAPI contrast and were pre-processed by CellPose’s “one-click image restoration”, and their segmentation required ad-hoc adjustments (merge and split of labels). To expedite processing times, masks were obtained from 2× downsampled images and then upscaled to original size. Spot-to-cell assignment was performed by iterating through each mask and running a 3D-maxima detection routine^36^ on the masked voxels for each color channel. Maxima with the same centroid coordinates on both channels were assumed to report lipofuscin and were discarded. Histograms of spot counts (Fig.4) were obtained from all the segmented cells in the field of view. The amplitude of Gaussian distributions overlaid on frequency data was adjusted so that the area under the curve (AUC) of both distributions matched. Summary expression was obtained by normalizing the spot counts associated with the labeled cell to the 10^th^ and 90^th^ percentile of the distribution obtained from all cells (≈1100) in the field of view (447×447×480µm). Other EASI-FISH quantifications were performed using the EASI-FISH pipeline^14^ via its Nextflow Tower interface (https://janeliascicomp.github.io/multifish/). Finalized charts were rendered in GraphPad Prism 10.3.1 (GraphPad Software).

### HortaCloud Implementation

The reconstruction software used locally by the MouseLight Project Team—Horta (*How Outstanding Researchers Trace Axons*)—was migrated to the cloud by adoption of Amazon’s AppStream Virtualized Desktop Infrastructure (VDI) framework, which allows the desktop software to be run in the cloud from a web browser by multiple concurrent users connected to the internet. This effort required several changes to the original codebase and is described in detail elsewhere^24^.

### Reconstructions

The detailed protocol is described in protocols.io^23^. In summary, cells with axonal arbors fully resolvable are identified and traced by an experienced annotator. Upon completion, a second annotator peruses the traced centerlines, annotating any mismatched branches. Differences are then extracted between the two annotations and a ‘best-guess’ consensus is obtained.

## Supporting information

morphoFISH Probes used

## Author Contributions

TAF, JC, PWT, and ME conceptualized the morphoFISH protocol. TAF, PWT, and ME established the morphoFISH protocol. MC prepared samples for imaging. TAF performed whole-brain imaging, image processing, and data analyses. ME performed EASI-FISH experiments. ET, MLL, MW, AKS, and JB provided reconstructions/brain registrations. DS, DJO, and KR developed the HortaCloud client. TAF and JTD defined the probing library. JTD designed the AAV2-retro experiment. NS, WK, JC, JTD, and TAF conceptualized the project. TAF oversaw all experiments. JTD and TAF wrote the manuscript with input from all authors. JTD and WK supervised the project. Remaining members of the MouseLight Project Team contributed to the general development of the project. The MouseLight Project Team during this time consisted of: Andrew Recknagel, Cameron Arshadi, Emily Tenshaw, Jayaram Chandrashekar, Mary Lay, Monet Weldon, Patrick Edson (Leap Scientific), and Tiago A. Ferreira. Joshua T. Dudman, Adam W. Hantman, Scott M Sternson, Karel Svoboda, Nelson Spruston, and Wyatt Korff were part of the MouseLight steering committee.

## Acknowledgments

The authors would like to thank Janelia Experimental Technologies, especially Alex Sohn, Daniel Flickinger, Jon Arnold, Mioara Gugiu, Steven Sawtelle, and Vasily Goncharov for hardware design, fabrication, and maintenance; Hyun Ah Yi and Janelia Virus Tools for viral reagents; Michael DeSantis for imaging assistance; Maya Tondravi and Gudrun Ihrke for coordinating experimental work; Janelia Scientific Computing, especially Adam Taylor, Ben Arthur, and David Ackerman for help with data processing; Ken Carlile and Scientific Computing Systems for help with data handling; and Janelia Vivarium, including Rae Demars and Sarah Lindo, for animal care and surgical assistance. We thank Luke Lavis for advice on morphoFISH chemistry and Sarah Plutkis for making Melphalan-X. Kari Close for EASI-FISH assistance. We thank Yin Liu and Geoffrey Meissner for critical reading of the manuscript. We are also grateful to the developers of the open source libraries used in this study and several laboratories/institutions for releasing data publicly, including the Allen Institute for Brain Science (Allen Mouse Common Coordinate Framework and Cell Types Database: RNA-Seq Data), and CEBSIT (ION) (fMOST reconstructions). This work was supported by the Howard Hughes Medical Institute.

This article is subject to HHMI’s Open Access to Publications policy. HHMI lab heads and project team leaders have previously granted a nonexclusive CC BY 4.0 license to the public and a sublicensable license to HHMI in their research articles. Pursuant to those licenses, the author-accepted manuscript of this article can be made freely available under a CC BY 4.0 license immediately upon publication.

## Data Availability

Neuron reconstructions were deposited at https://ml-neuronbrowser.janelia.org/. Imagery is hosted on AWS Open Data (https://registry.opendata.aws/janelia-mouselight/) and accessible in all HortaCloud instances https://hortacloud.org/. Scripts used in this manuscript will be available at https://github.com/MouseLightPipeline/morphoFISH.

---

**Figure S1 (Related to Fig. 3). Inspection of local neuropil is insufficient to resolve the majority of cross-over errors.**

**a–i**: Exemplars of cross-over locations between axonal branches (across different cells and samples). For each panel, original image is shown on top and resolved skeletonized reconstructions at the bottom. Neurites associated with the same cell share the same color. Scale bars are indicated.

**j**: Amount of traced cable length (mm) required to resolve each location. Typically, large distances required tracing across multiple neurites and/or cells. All locations required tracing beyond the displayed insets. MouseLight IDs of complete neurons involved in each exemplar: A,C: AA1566; E: AL0002; F: AA1596; G: AA1596, AA1507; H: AA1564, AA1565; I: AA1559, AA1560, AA1571.

**Figure S2 (Related to Fig. 3). Fluorophore availability is not the exclusive bottleneck to reconstruction completeness between neighboring cells.**

**a.** Examples of cortical (CTX), hippocampal (HPF), and thalamic (TH) neurons. Endogenous GFP levels color-coded with normalized voxel intensities mapped to a warmer hue. Cells that could not be traced to completion are marked as “Partial”, while cells traced successfully by two independent annotators are marked as “Complete”. Scale bar: 20µm.

**b.** Linear regression between amount of traced cable and normalized soma intensity at the soma. Dashed lines indicate 95% confidence intervals. N= 39 cells, R^2^ = 0.1096, Pearson r = 0.3311.

**c.** Normalized soma intensities across the 3 groups. Median (full line) and quartiles (dashed lines) are indicated. N (*Partial*/*Complete*): 11/23 (CTX), 2/10 (HPF), 11/7 (TH).

Data gathered from MouseLight Sample ID 2018-08-01 available in HortaCloud.

**Figure S3 (Related to Fig. 4). Expression pattern of dual-stereotaxic injection revealing STR^IPSI^/PF^CONTRA^ GFP^+^/tdTom^+^ cells in the ACA (overview).**

Montage depicts anterior-posterior (A-P) sequence of contiguous 100µm-thick coronal sections. Fluorophores and scale bar are indicated.

**Figure S4 (Related to Fig. 4, S3). Expression pattern of dual-stereotaxic injection revealing STR^IPSI^/PF^CONTRA^ GFP^+^/tdTom^+^ cells in the ACA (detailed insets).**

Magnified inserts from the region highlighted in Figure S3.

## Supplemental information

**Table S1. Summary of probe library for sampling diversity of neuronal types in the neocortex.** Data gathered from public datasets of RNAseq data^12,43^ that have identified 13 groups of excitatory (i.e., Slc17a7-expressing) subtypes in the mouse prefrontal cortex (rows). Columns list genes selected as cluster “leads”, chosen for their discriminatory power and abundance, so that—typically—only 3 to 4 genes are required to distinguish subtypes within a cortical layer. Color coding reflects binarized expression.

**Data S1. List of HCR probes used in this study** (XLS file).

