## Supplementary figures and images for "Discovery of neuronal cell types by pairing whole cell reconstructions with RNA expression profiles"

### FigS1

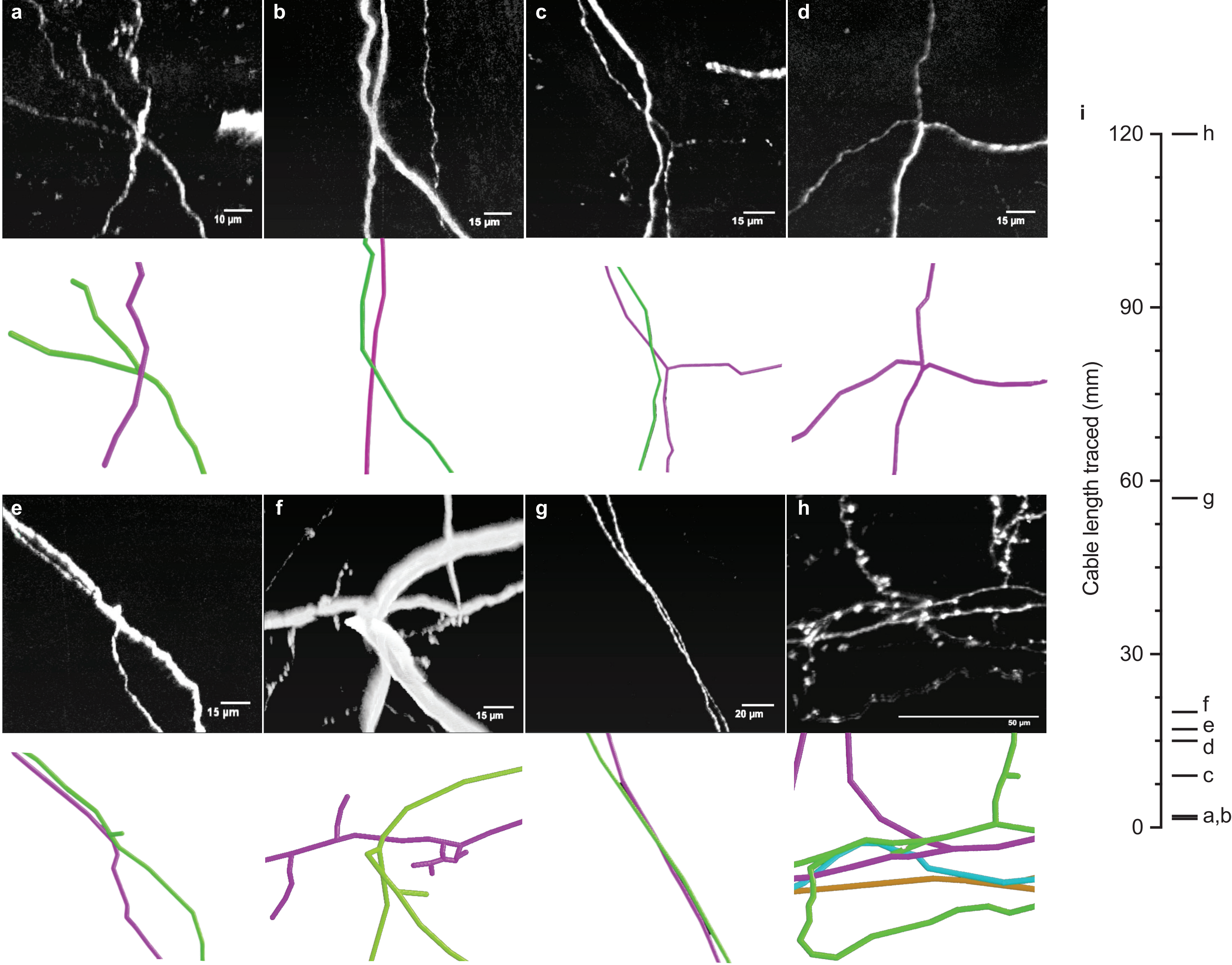


Figure S1 | Ferreira et al.

### FigS2

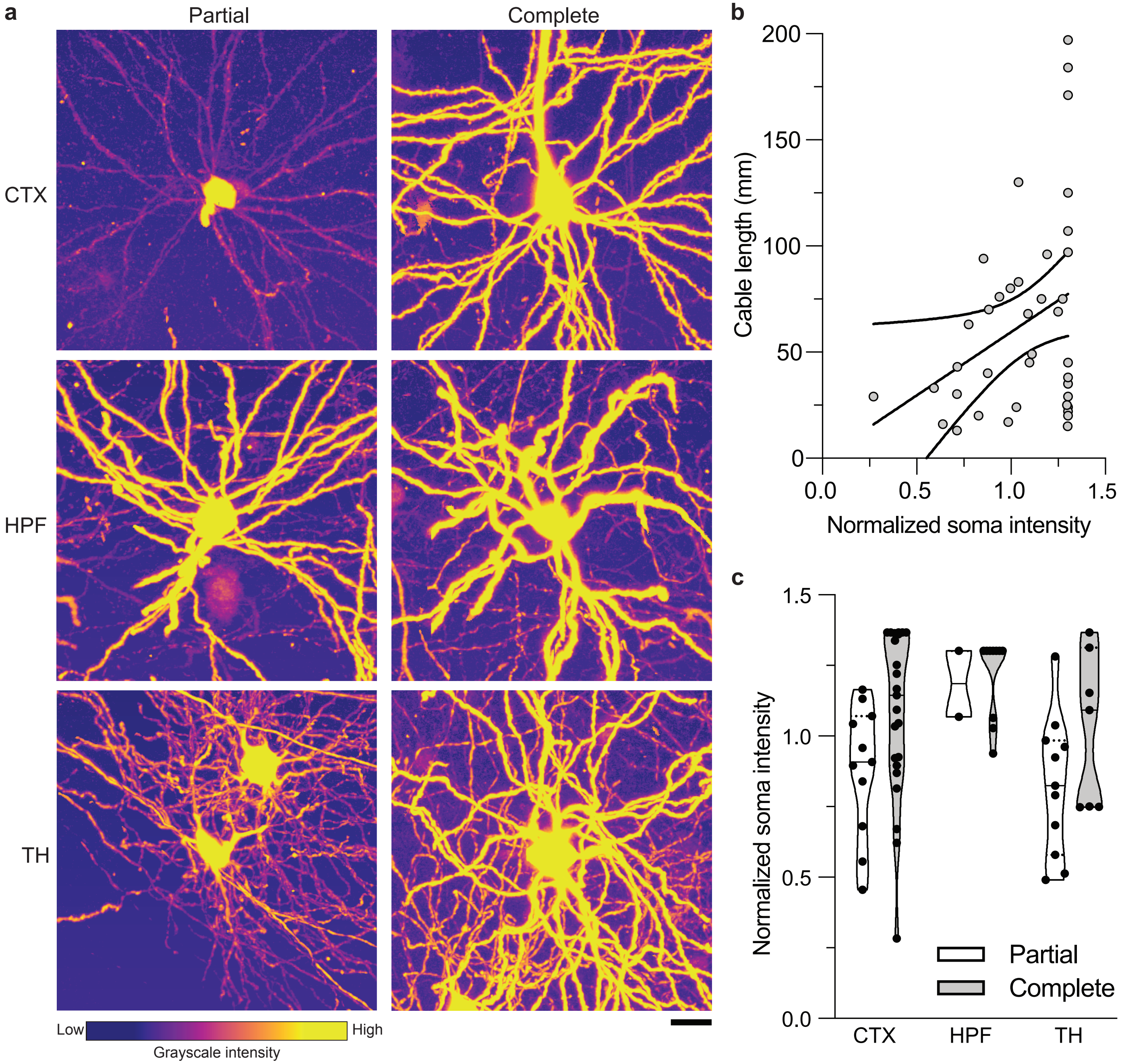


Figure S2 | Ferreira et al.

### FigS3

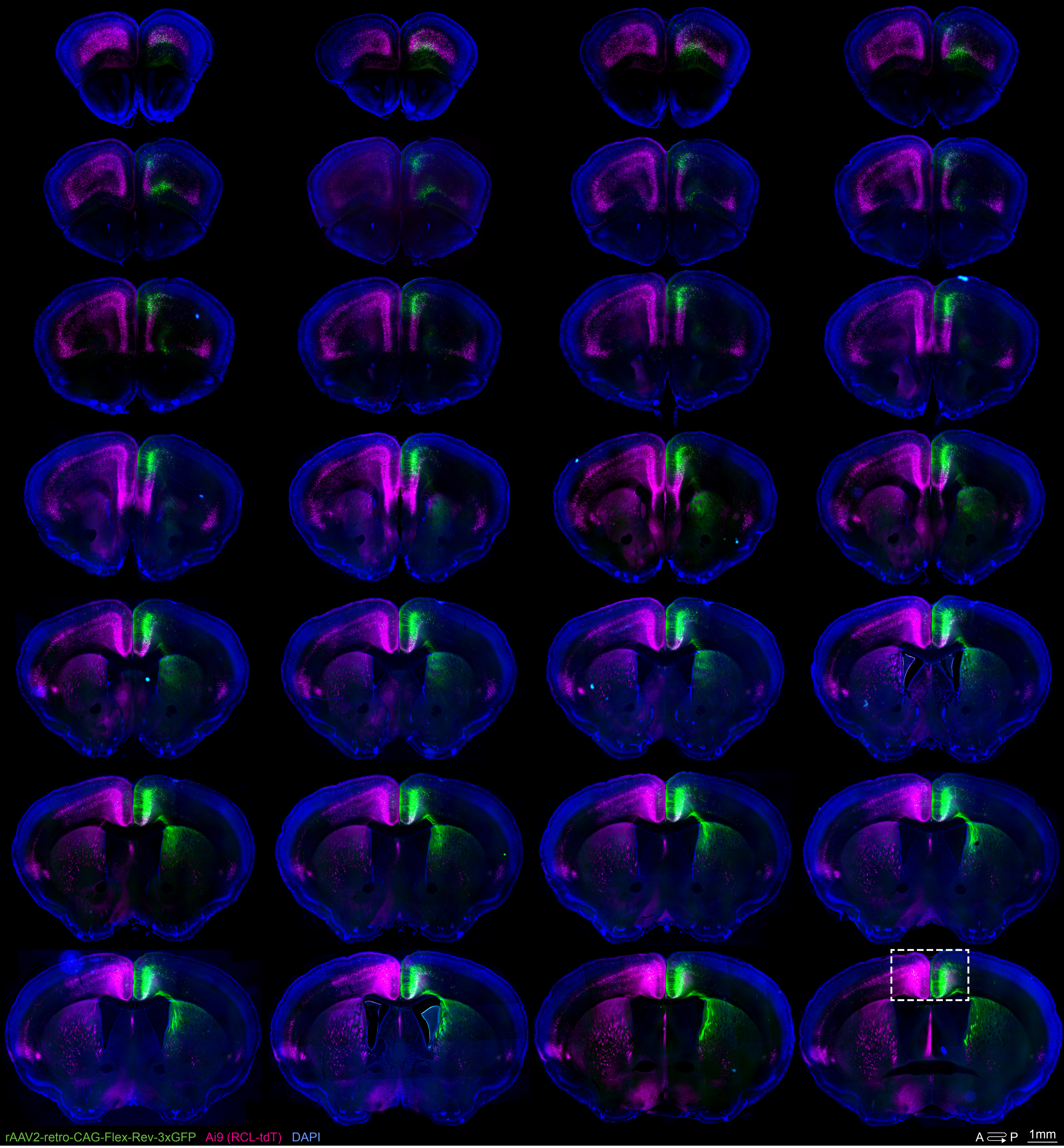


Figure S3 | Ferreira TA et al.

### FigS4

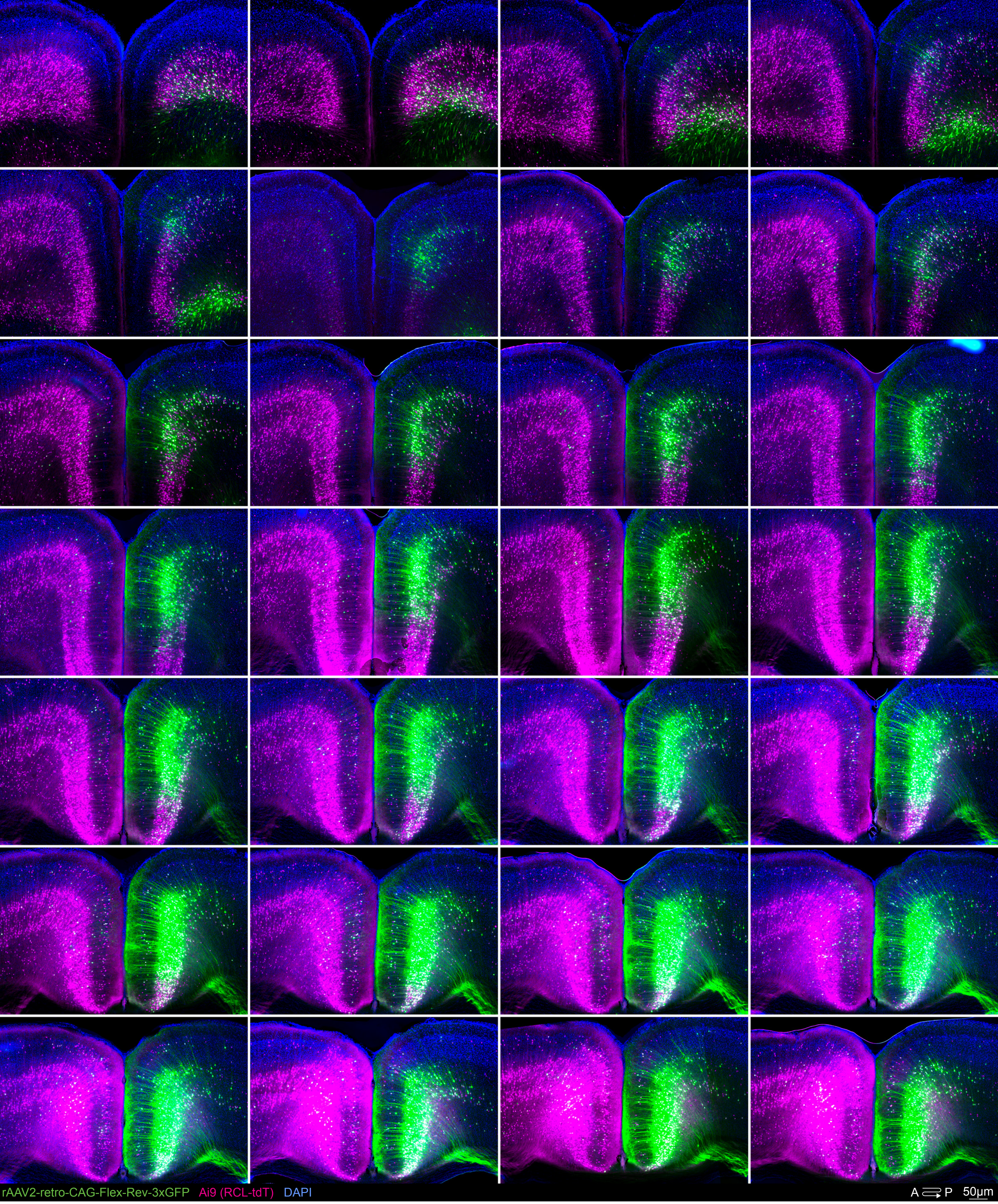


Figure S4 | Ferreira et al.

### Table S1

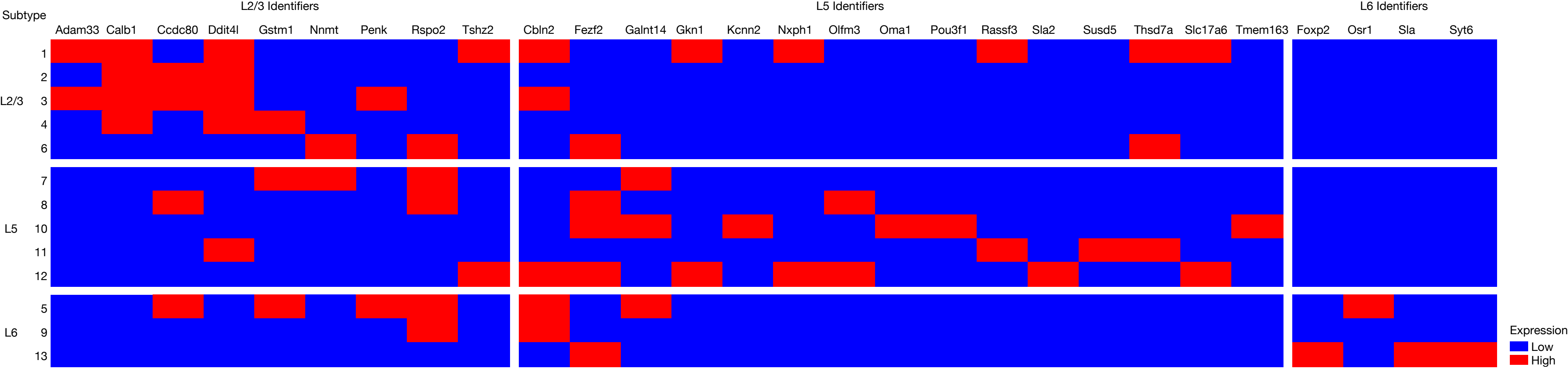
